# *Staphylococcus aureus* enhances influenza A virus replication through modulation of host cell vesicular transport

**DOI:** 10.64898/2026.10.12.758568

**Authors:** Veerle Appels, Karen Nickel, Elise Therrien, Isabella Schmidt, Lily Sheridan, Megan VanderWal, Rachel Witt, Mariya I. Goncheva

## Abstract

Influenza A virus (IAV) is a major respiratory pathogen that causes substantial morbidity and mortality worldwide. A significant complication of primary IAV infection is secondary bacterial co-infection. Here, we report that *Staphylococcus aureus* enhances IAV replication through the activity of the surface protein fibronectin-binding protein A (FnbA). The pro-viral activity of FnbA is conserved across multiple IAV strains and is associated with reduced abundance of the host vesicular trafficking proteins VPS26A and VTA1. FnbA does not substantially increase viral protein abundance or alter intracellular distribution. Instead, pro-viral activity is associated with a greater proportion of infectious progeny particles and increased release of infectious virions from infected cells. Thus, we identify FnbA as a bacterial factor that enhances IAV replication and demonstrate that *S. aureus* can manipulate host cell processes to promote the production and release of infectious virus.

## Introduction

Influenza A virus (IAV) is an enveloped, negative-sense RNA virus with a segmented genome that causes substantial respiratory disease worldwide. Seasonal IAV epidemics are estimated to cause 290,000–600,000 deaths annually, while the emergence of antigenically distinct strains can result in pandemics associated with increased morbidity and mortality^1–3^. A significant complication of primary IAV infection is secondary bacterial co-infection. Historical analyses of the 1918 H1N1 pandemic identified bacterial involvement in 70–90% of autopsy cases^3^, and despite the widespread availability of antibiotics, co-infection remained associated with substantial mortality during the 2009 influenza pandemic^4,5^. *Staphylococcus aureus* is currently the most frequently identified bacterial pathogen in influenza-associated co-infections^6–10^.

Much of our understanding of IAV–*S. aureus* interactions has focused on how viral infection increases susceptibility to subsequent bacterial disease. IAV disrupts the respiratory epithelium and alters the pulmonary environment, creating conditions that favour bacterial colonization and growth^11–14^. Concurrent changes in innate and adaptive immune responses also impair bacterial recognition and decrease bacterial clearance^11,12,15–17^. Consequently, IAV is generally viewed as the primary driver of the interaction, with subsequent bacterial disease arising from virus-induced changes in the host.

Considerably less is known about the reciprocal effects of *S. aureus* on IAV replication. Only three bacterial factors have been implicated in this process. The *S. aureus* exoprotein expression regulatory system (SaeRS) contributes to pathogenesis during IAV–*S. aureus* co-infection in mice^18^, while phenol-soluble modulins (PSMs) increase cytotoxicity in IAV-infected lung epithelial cells^19^. The secreted *S. aureus* Lipase 1 has also been shown to enhance IAV replication in primary cells by modifying host lipids and increasing infectious virus production^20^. These findings demonstrate that *S. aureus* can influence the course of viral infection, but the bacterial factors and host processes that contribute to these effects remain to be fully characterized.

The capacity of *S. aureus* to modify the cellular environment is particularly relevant to IAV, which relies extensively on host machinery throughout its replication cycle. Vesicular trafficking contributes to multiple stages of IAV infection. The retromer complex, comprising vacuolar protein sorting-associated proteins 26A (VPS26A), 29 (VPS29), and 35 (VPS35), controls endosomal cargo sorting and recycling^21^ and has been implicated in IAV entry^22^. Vesicle trafficking 1 (VTA1) functions as a cofactor for vacuolar protein sorting-associated protein 4 (VPS4), promoting VPS4 activity and disassembly of the endosomal sorting complexes required for transport III (ESCRT-III) machinery^21,23^. IAV does not require VPS4 or the canonical ESCRT machinery for efficient budding and release^24,25^. However, its broader dependence on host trafficking provides multiple opportunities for perturbations in these pathways to influence viral replication.

Here, we developed an *in vitro* co-infection model to investigate the molecular interactions between the host, IAV, and *S. aureus*. Using systems approaches combining proteomic analysis with *S. aureus* genome-wide genetic screening, we identified the surface protein fibronectin-binding protein A (FnbA) as a bacterial determinant that enhances IAV replication. The pro-viral activity of FnbA is associated with reduced abundance of the host trafficking proteins VPS26A and VTA1. FnbA-dependent enhancement occurred late in the viral replication cycle, increasing the proportion of infectious progeny particles and promoting their release from infected cells. Together, these findings identify a previously unrecognized function of FnbA in modulating host cell biology and demonstrate that this activity can promote IAV replication during co-infection.

## Materials and methods

### Cell lines and culture conditions

Human alveolar epithelial A549 cells (A549; ATCC CCL-185) and Madin-Darby canine kidney (MDCK; ATCC CCL-34) cells were maintained at 37°C, 5% CO_2_. A549 cells were cultured in Dulbecco’s Modified Eagle Medium (DMEM) supplemented with 10% (v/v) fetal bovine serum (FBS). MDCK cells were cultured in DMEM supplemented with 5% (v/v) FBS and 2 mM L-glutamine.

### Viruses

Influenza A virus (IAV) strains used in this study are listed in Table 1. IAV stocks were propagated in confluent MDCK monolayers. Cells were infected at a multiplicity of infection (MOI) of 0.01 for 1 hour at 37°C and 5% CO_2_. The inoculum was then removed and replaced with virus growth medium consisting of DMEM with 2 mM L-glutamine, 0.14% (w/v) bovine serum albumin (BSA), and 1 μg/mL N-Tosyl-L-phenylalanine chloromethyl ketone (TPCK) treated trypsin. Cultures were maintained until extensive cytopathic effect was observed (typically 48-96 h post-infection (hpi)). Supernatants were clarified by centrifugation, aliquoted, and stored at −80°C. Infectious virus titres were determined by plaque assay on MDCK cells, under an Avicel overlay^26^.

**Table 1.** A list of viruses used.

| Virus Strain | Abbreviation | Source |
| --- | --- | --- |
| A/Puerto Rico/8/1934 (H1N1) | PR8 | BEI |
| A/Wilson-Smith/1933 (H1N1) | WSN | BEI |
| A/WSN/33-PA-2A-NLuc (H1N1) | WSN-NanoLuc | <sup>27</sup> |
| A/Baltimore/JH-22377/2022 (H1N1) (seasonal isolate) | sH1N1 | BEI |
| A/Baltimore/JH-335/2022 (H3N2) (seasonal isolate) | sH3N2 | BEI |

### Bacterial growth conditions

Bacterial strains and plasmids used in this study are listed in Supplementary Table 1. *Escherichia coli* was grown in Luria Broth (LB) and *Staphylococcus aureus* was grown in tryptic soy broth (TSB) at 37 °C, with shaking at 200 rpm, unless otherwise stated. Where appropriate, media were supplemented with erythromycin (3 µg/mL), chloramphenicol (12 µg/mL) or ampicillin (100 µg/mL). Solid media were supplemented with 1.5% (w/v) Bacto agar.

*S. aureus* strain USA300 LAC, cured of the 27-kb plasmid conferring antibiotic resistance, was used as the wild-type (WT) strain for mutant generation, unless otherwise stated. For complementation, the full-length genes were amplified by polymerase chain reaction (PCR) using primers listed in Supplementary Table 2, ligated into pALC2073^28^ and recombinant plasmids transformed into *E. coli*. Plasmids were then passaged through *S. aureus* RN4220^29^, and subsequently transformed into the strain of interest.

### Viral-bacterial coinfections

Co-infections were established in confluent A549 monolayers in 12 well plates. Cells were infected with IAV at an MOI of 3 in a total volume of 100µl per well for 1 h at 37°C and 5% CO_2_. The inoculum was then removed, and cells were washed once with phosphate buffered saline (PBS). Extracellular virus was removed by treatment with acid wash buffer (10 mM NaCl, 0.1% HCl, pH 3.0) for 90 sec. Cells were then washed three times with PBS and fresh DMEM was added. *S. aureus* from overnight cultures were washed twice with DMEM, normalized to OD_600_ equivalent to 2×10^7^ colony forming units (CFU)/mL and 5µl were added per well to virus infected wells to achieve the desired final concentration of 1×10^5^ CFU/mL. Mock-infected, virus-only, and bacteria-only controls were included in all experiments. Cells were incubated for the indicated times following bacterial addition.

At harvest, supernatants were collected for pathogen quantification. Bacterial burdens were determined by serial dilution in PBS and plating on TSA. Remaining supernatants were centrifuged at 17,000 × g for 3 min to remove bacteria and cellular debris, filtered through 0.22 μm polyethersulfone (PES) filter and stored at −80°C. Following supernatant removal, cell monolayers were washed three times with PBS and lysed in radioimmunoprecipitation assay buffer (RIPA) buffer supplemented with protease inhibitors. Lysates were incubated at 4°C for 20 min, then clarified by centrifugation at 12,000 × g for 10 min, and the resulting supernatants were stored at −20°C until further analysis.

### Luciferase screening

The Nebraska transposon mutant library^30^ was maintained as frozen stocks in 96-well plates. For screening experiments, frozen stocks were inoculated into fresh TSB in 96-well plates and cultured overnight at 37°C with shaking at 180 rpm. Overnight cultures were diluted 40-fold in DMEM, and the resulting bacterial suspensions were used (see below). A549 cells were grown to confluence in 96-well plates and infected with IAV-NanoLuc at an MOI of 2 in 50 μL DMEM per well. Following a 1 h incubation at 37°C and 5% CO_2_, the viral inoculum was removed, cells were washed once with PBS, and 200 μL fresh DMEM was added to each well. Transposon mutant cultures were added at 5 µl per well. At 12 hpi, luciferase activity was quantified using the Nano-Glo Luciferase Assay System according to the manufacturer’s instructions.

### Quantitative proteomics

For proteomic analysis, coinfections were established as described above but in T25 flasks and harvested at 6 h post bacterial addition. Following supernatant removal, cell monolayers were washed three times with PBS, scraped into suspension, and collected by centrifugation. Pellets were frozen at −80°C and subsequently analyzed by Liquid Chromatography-Tandem Mass Spectrometry (LC-MS/MS) at the UVic-Genome BC Proteomics Centre.

Cell pellets were lysed, digested with trypsin, and analyzed by reverse-phase LC-MS/MS using an Orbitrap Fusion Tribrid mass spectrometer. MS/MS spectra were acquired using data-dependent acquisition and searched using FragPipe with MSFragger against a custom protein database containing *Homo sapiens*, *Staphylococcus aureus* USA300, and influenza A virus (A/Puerto Rico/8/1934 H1N1) sequences obtained from UniProt. Carbamidomethylation of cysteine was specified as a fixed modification, while methionine oxidation and protein N-terminal acetylation were specified as variable modifications.

### Western blotting

Protein lysates were separated by Sodium Dodecyl Sulfate-Polyacrylamide Gel Electrophoresis (SDS-PAGE) and transferred to nitrocellulose membranes. Membranes were blocked in PBS containing 7% (w/v) skim milk powder, 10% (v/v) calf serum, and 10% (v/v) horse serum and probed with antibodies against β-actin (Fisher AM4302, 1:1500), VTA1 (Invitrogen PIPA521831, 1:750), VPS26A (Abcam ab23892, 1:1500), or influenza A virus nucleoprotein (NP) (Abcam ab20343, 1:1500). Following incubation with IRDye-conjugated secondary antibodies (1:10,000), membranes were imaged using a LI-COR Odyssey imaging system.

### Immunofluorescence

A549 cells were grown to confluence on 18 mm glass coverslips. To synchronize viral entry, a cold-sync infection was performed as previously described^31,32^ with minor modifications. Briefly, cells were washed twice with ice-cold PBS, incubated on ice for 15 min, and chilled virus inoculum (MOI = 6) was added at 4°C for 60 min to allow viral attachment. The inoculum was then removed, cells washed twice with cold PBS, pre-warmed DMEM was added and cells were shifted to 37°C for 30 min. Following this incubation, media was removed, cells washed twice with pre-warmed PBS and fresh pre-warmed DMEM added. *S. aureus* to a final concentration of 1 × 10^5^ CFU/mL was added, and this was designated as 0 hpi. At indicated times post infection, supernatant was removed and cells were fixed with 4% paraformaldehyde for 20 min at room temperature (RT). Cells were then washed 3 times with PBS and permeabilized with 0.1% (v/v) Triton X-100 for 5 min. Samples were blocked for 4-6h at RT with 10% (w/v) BSA + 10%(v/v) guinea pig serum + 10µg/mL of purified guinea pig IgG. This specific blocking cocktail was optimized for maximum decrease in the non-specific binding of IgG that is mediated by *S. aureus* protein A and second immunoglobulin binding protein (Sbi). Following blocking, cells were incubated with primary antibodies in blocking buffer overnight at 4°C. Antibodies used were VPS26A (Abcam ab23892, 1:500) and IAV nucleoprotein (NP) (Abcam ab20343, 1:500). Coverslips were washed three times with PBS before incubation with species-specific Alexa Fluor-conjugated secondary antibodies for 1 h at RT. Following three PBS washes, nuclei were counterstained with Hoechst (1:7500) for 20 min at RT. Coverslips were washed with PBS and mounted using ProLong Diamond Antifade Mountant.

### Imaging

Widefield fluorescence images were acquired using a Nikon Eclipse Ti2-E inverted microscope equipped with a Plan Apo λD 60×/1.42 NA oil immersion objective and a Teledyne Photometrics Kinetix sCMOS camera. Images were acquired using NIS-Elements AR version 6.02.03 (Nikon). All samples within an experiment were imaged using identical acquisition settings and processed identically. All images were deconvolved using the Rich-Lucey method and converted to maximum-intensity projections prior to analysis. Individual cells were selected from each field of view, and the mean fluorescence intensity of the host and viral proteins was quantified for each cell.

To assess the spatial distribution of fluorescence, four line profiles were generated across each selected cell. To account for differences in cell size, each line profile was divided into six equal sections spanning the cell region outside of the nucleus. Fluorescence within each section was quantified by calculating the area under the curve of the intensity profile. Each line profile was then assigned a score from 1 to 6 corresponding to the section with the highest signal from area under the curve. Each line profile was scored only once, resulting in four independent spatial measurements per cell.

### Particle to PFU ratio

Co-infections were established following synchronization of viral entry as described above for immunofluorescence. Viral RNA was extracted from virus-containing supernatants and reverse transcribed using SuperScript IV Reverse Transcriptase (Thermo Fisher Scientific) with universal IAV Uni12 primer (Supplementary Table 2). Viral genome copy number was determined by quantitative PCR targeting the influenza A virus M gene using absolute quantification against a plasmid standard curve (10^7^–10^1^ copies per reaction), as previously described^33,34^ with minor modifications using primers listed in Supplementary Table 2. Infectious virus titres were determined by standard plaque assay as described above. To allow for accurate quantification of all virions, these samples were not filtered at harvest times. Instead, plaque assays were performed in the presence of 150µg/mL of gentamicin in the overlay media, to kill any remaining bacteria. Particle-to-PFU ratios were calculated by dividing viral genome copies per mL by the corresponding infectious titre (PFU/mL) from the same virus sample.

### Cell-associated and released virus assay

To quantify cell-associated and released infectious virus, at the indicated time post-infection, cells were scraped into the culture medium and centrifuged at 1,600 × *g* for 5 min to separate the cell pellet from the supernatant. The resulting supernatant was collected for quantification of released infectious virus. Cell pellets were subjected to three freeze-thaw cycles to recover cell-associated virus. Infectious virus in both fractions was quantified by plaque assay. To allow for accurate quantification of all virions, these samples were not filtered at harvest times. Instead, plaque assays were performed in the presence of 150µg/mL of gentamicin in the overlay media, to kill any remaining bacteria.

### Statistical analysis

Statistical analyses were performed using GraphPad Prism. Data are presented as mean ± standard error of the mean (SEM). Statistical tests used for individual experiments are indicated in the corresponding figure legends. Differences were considered statistically significant at p≤0.05.

## Results

### Staphylococcus aureus enhances IAV replication in vitro

To investigate interactions between IAV, *S. aureus*, and host cells in the absence of an immune system, we established an *in vitro* co-infection model (Fig. 1A). Preliminary experiments showed substantial *S. aureus*-induced cytotoxicity by ∼13 hours post infection (hpi) (data not shown). Therefore, all subsequent analyses were performed no later than 12 hpi, allowing for a complete IAV replication cycle to occur before the onset of detectable cytotoxicity.

**Figure 1.**
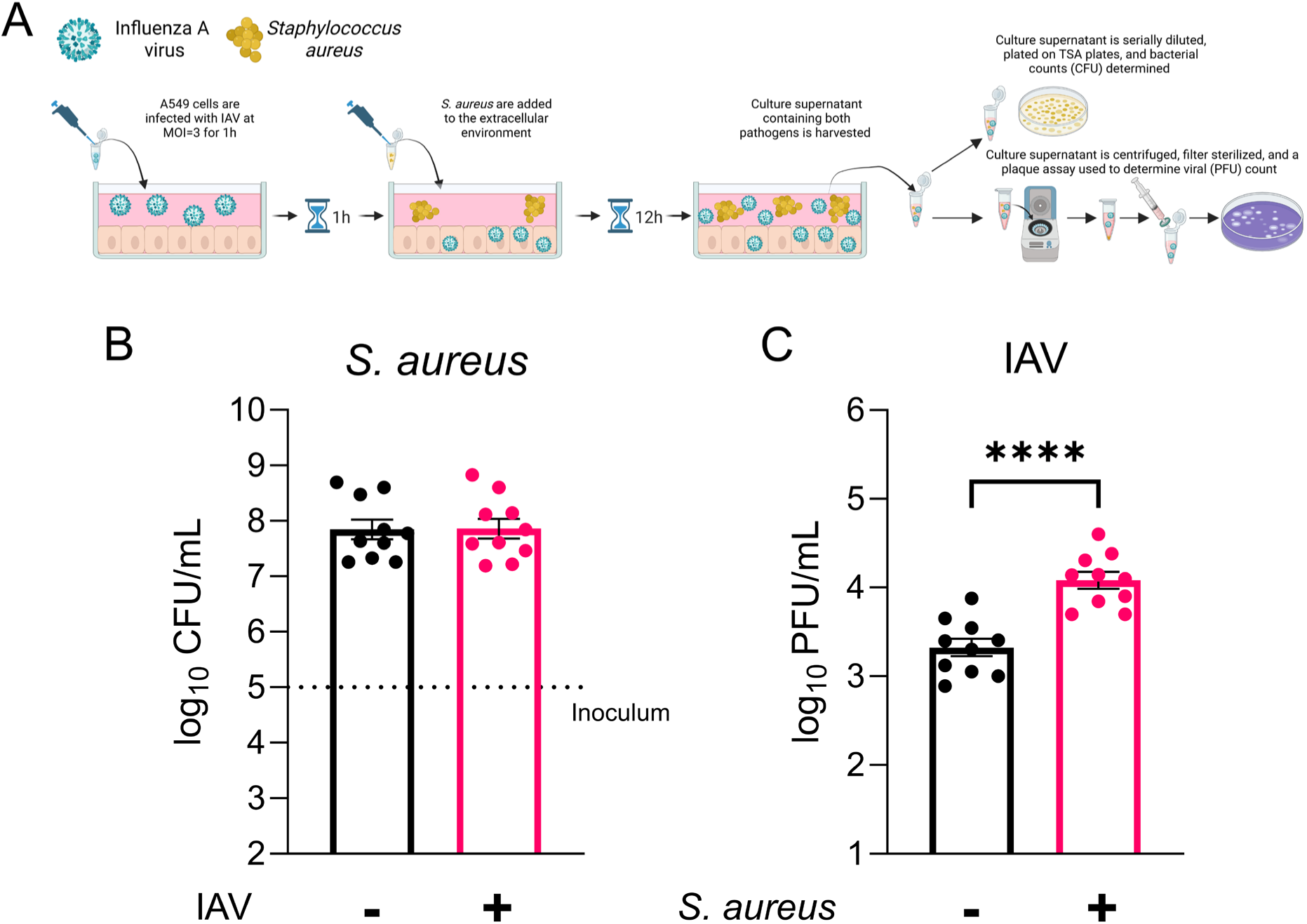
***S. aureus* enhances IAV replication. A)** Schematic of the infection model. A549 cells in a 12 well plate were infected with A/Puerto Rico/8/34 at MOI 3 for 1h. Inoculum was removed, extracellular virus lysed with an acid wash for 90 sec, and fresh media added. 1×10^5^ CFU/mL of *S. aureus* were added to the extracellular environment. **B)** At 12h post infection (hpi) a sample from the supernatant was taken, serially diluted in PBS, plated on TSA plates and incubated at 37°C for 18h. The dotted inoculum line represents the number of bacteria added at 0h. **C)** Samples from **B)** were collected, centrifuged at 17,000 x *g* for 3 minutes and the resulting supernatant passed through a 13 mm, 0.22 µm filter. A standard plaque assay was then performed on MDCK cells to determine infectious virus titre. Data shown are mean ± SEM of 10 experiments. **** is p value ≤ 0.0001, based on a two-tailed, paired student’s t test.

To characterize this model, we first quantified bacterial and viral replication. *S. aureus* replicated throughout the course of infection, with no difference between mock- and IAV-infected cells (Fig. 1B). In contrast, infectious IAV titres increased approximately 10-fold in the presence of *S. aureus* (Fig. 1C). To determine whether this phenotype extended beyond *S. aureus* USA300, we repeated infections using multiple *S. aureus* strains. Although bacterial replication varied, IAV infection did not affect bacterial growth (Supplementary Fig. 1A). Apart from Newman, all tested strains enhanced IAV replication to similar levels (Supplementary Fig. 1B).

Together, these findings establish a robust *in vitro* model for investigating IAV-*S. aureus* interactions and demonstrate that co-infection with *S. aureus* enhances the production of infectious IAV particles.

### Co-infection of IAV and *S. aureus* induces changes in the abundance of host cell trafficking proteins

Given that *S. aureus* enhances IAV replication, we hypothesized that the bacterium induces changes in host cells that promote infectious virus production. To identify these changes, cells were infected as in Fig. 1A and harvested at 6 hpi for label-free proteomic analysis. Analysis of differentially abundant proteins revealed 34 proteins that differed significantly across the experimental groups (Fig. 2A). Direct comparison of IAV-infected and IAV-*S. aureus* co-infected cells identified three candidates (Fig. 2B). These were autophagy-related protein 9A (ATG9A), vacuolar trafficking-associated protein 1 (VTA1; also known as LIP5), and vacuolar protein sorting protein 26A (VPS26A). ATG9A abundance was altered in co-infected cells, but this change was not specific to the comparison between IAV-infected and IAV-*S. aureus* co-infected cells (Fig. 2A). Therefore, this protein was not investigated further.

**Figure 2.**
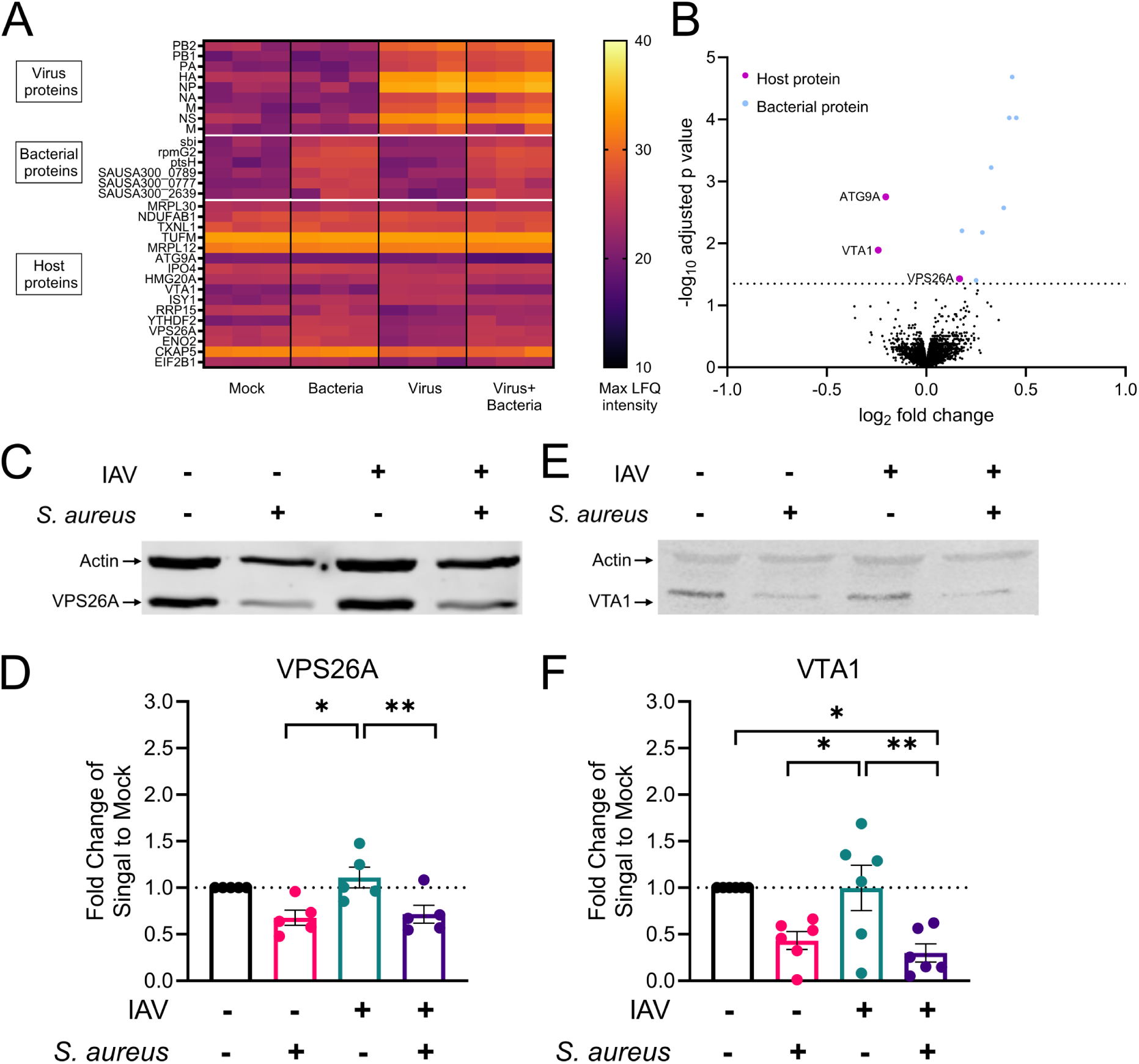
***S. aureus* co-infection triggers changes in the abundance of host cell trafficking proteins.** Cells were infected as in Figure 1A. At 6hpi cells were harvested and label-free proteomics performed. **A)** Heat map of differentially abundant proteins between all groups. Each column represents a distinct biological sample. **B)** Volcano plot comparing virus to virus with bacteria samples. Cells were infected as in Figure 1A. At 12hpi cells were harvested, lysed and western blot performed for **C)** VPS26A and **E)** VTA1. Cellular actin was measured concurrently in each sample. **D)** and **F)** Analysis of western blot images was performed. For each image, values were normalized to actin and are shown as fold change of mock. For statistical analysis, actin normalized values were converted to log2, and a one-way ANOVA with repeated measures was performed, with Turkey’s multiple comparison correction. ± SEM of 4-6 experiments. * is p value ≤ 0.05, ** is p value ≤ 0.01.

To validate the changes identified by proteomic analysis, VTA1 and VPS26A abundance was assessed by Western blot. At 6 hpi, neither protein differed between infection conditions (Supplementary Fig. 2). By 12 hpi, *S. aureus* reduced VTA1 and VPS26A abundance by approximately 50% in both bacteria-only and co-infected cells, while IAV infection alone had no effect (Fig. 2C-F). Together, these data demonstrate that *S. aureus* alters the levels of host vesicle trafficking proteins, including during IAV-*S. aureus* co-infection.

### *S. aureus* enhances IAV replication via the SaeRS two-component system

As the pro-viral effect observed in Fig. 1 varied among the *S. aureus* strains tested (Supplementary Fig. 1), we hypothesized that this phenotype is mediated by specific bacterial factors rather than the presence of *S. aureus* itself. To test this, we adapted our co-infection model for high-throughput analysis. We used IAV encoding Nano luciferase (IAV-Nano-Luc), where luminescence levels provide a measure of viral replication^35^. Virus replication was first assessed with this reporter system in the presence or absence of *S. aureus.* Uninfected and bacteria-only samples showed only background signal levels, while IAV infection resulted in a ∼100-fold increase in signal (Fig 3A). Importantly, addition of *S. aureus* resulted in a ∼15-fold increase in luminescence compared with IAV alone, recapitulating the pro-viral phenotype observed in our original co-infection model (Fig 3A).

**Figure 3.**
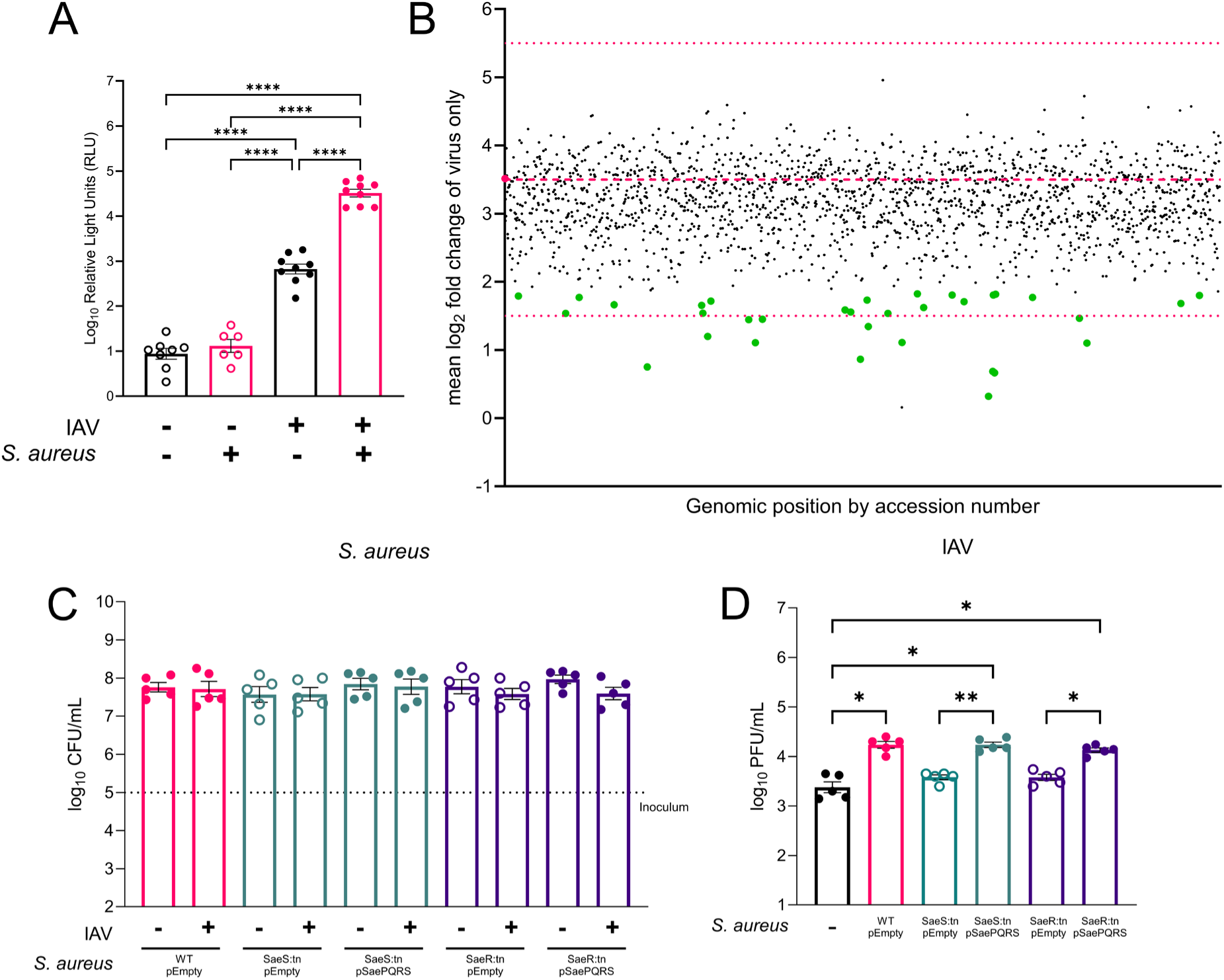
A genome wide screen of non-essential *S. aureus* genes identified that SaeRS contributes to pro-viral activity. **A)** A549 cells in a 96 well plate were infected with IAV WSN-Luc at a MOI of 2 for 1h. Inoculum was removed and serum-free media added. 1×10^5^ CFU of *S. aureus* were added to the media. At 12h post infection (hpi) a sample from the supernatant was taken, mixed with an equal volume of Nano-Glo reagent and luminescence measured on a plate reader. Data shown are mean ± SEM of 6 experiments. **** is p value ≤ 0.0001, based on a one-way ANOVA. **B)** Cells were infected as in **A).** ∼5×10^4^ CFU of WT or mutant *S. aureus* (total of 1920 individual mutants) bacteria were added to the media after virus infection. At 12hpi a sample from the supernatant was taken, mixed with equal volume of Nano-Glo reagent and luminescence measured on a plate reader. All data were normalized to the virus only levels for each experiment and converted to log2 fold change. The mean value from 4 independent experiments is plotted. The dark dotted line represents the mean fold change of the virus + WT *S. aureus* sample. Thin dotted lines represent the boundaries of −2 and +2 log_2_ fold change from the WT value. Green values were identified as significant, with p value ≤ 0.05, when compared to the virus + WT *S. aureus* sample, based on a one-way ANOVA, with Fisher’s LSD test. **C)** Cells were infected as in Figure 1A, with WT or mutant *S. aureus*. At 12hpi a sample from the supernatant was taken, serially diluted in PBS, plated on TSA plates and incubated at 37°C for 18h to determine CFU/mL. **D)** Samples from **C)** were taken, centrifuged at 21,000 x *g* for 3 minutes and the resulting supernatant passed through a 13 mm, 0.22 µm filter. A standard plaque assay was then performed on MDCK cells. Data shown are mean ± SEM of 4-9 experiments. * is p value ≤ 0.05, ** is p value ≤ 0.01, based on a repeated measures one-way ANOVA with Turkey’s multiple comparison correction.

Having validated the IAV-Nano-Luc system, we performed a genome-wide screen of non-essential *S. aureus* genes using an ordered library of transposon insertion mutants^30^. A549 cells were infected with IAV-Nano-Luc, followed by addition of 1920 individual *S. aureus* mutants. At 12 hpi, luminescence was measured and normalized to IAV-only controls. This screen identified 33 *S. aureus* mutants with significantly altered pro-viral activity (Fig. 3B, Supplementary Table 3). Nine mutants showed altered growth in the presence of A549 cells (Supplementary Fig 3) and were excluded from further analysis. Of the remaining 24 candidates, 15 were previously characterised genes.

Among the mutants of interest were components of the SaeRS two-component regulatory system^36^. SaeRS has previously been implicated in IAV–*S. aureus* co-infection^18^, and strain Newman, which lacked pro-viral activity in our model (Supplementary Fig. 1), carries a known mutation in this system^37,38^. We therefore selected SaeRS for additional analysis. To investigate this further, we generated complemented strains carrying the *saePQRS* operon *in trans*. Cells were infected as in Fig. 1A using WT, *saeR* or *saeS* mutants, or complemented strains. Bacterial replication did not differ between groups (Fig. 3C). Mutation of either *saeS* or *saeR* eliminated pro-viral activity, while complementation with the *saePQRS* operon restored the phenotype to WT levels (Fig. 3D). Taken together, these data demonstrate that the SaeRS two-component system is a major contributor to the *S. aureus* mediated increase in IAV replication.

### *S. aureus* fibronectin binding protein A increases IAV replication through changes in VPS26A and VTA1 abundance

Given the intracellular localization of SaeR and membrane localization of SaeS^36^, we hypothesized that the pro-viral phenotype is mediated by one or more proteins regulated by this two-component system rather than through a direct interaction with the host cell. To identify potential effectors, we used the IAV-Nano-Luc assay to screen the 33 SaeRS-regulated genes present in the *S. aureus* USA300 genome^36^. Of these, mutation of SAUSA300_2441, encoding fibronectin-binding protein A (FnbA), significantly reduced pro-viral activity (Supplementary Fig. 4). To validate the contribution of *fnbA*, we used a previously established Δ*fnbAB* mutant, in which both *fnbA* and its functional homologue *fnbB*^39^ are deleted, together with a complemented strain carrying *fnbA* in trans^40^. Using our co-infection model (Fig. 1A), we saw bacterial replication was comparable between WT, Δ*fnbAB*, and complemented strains (Supplementary Fig 5). In contrast, deletion of *fnbAB* eliminated the pro-viral effect, while complementation with *fnbA* restored the phenotype (Fig. 4A).

**Figure 4.**
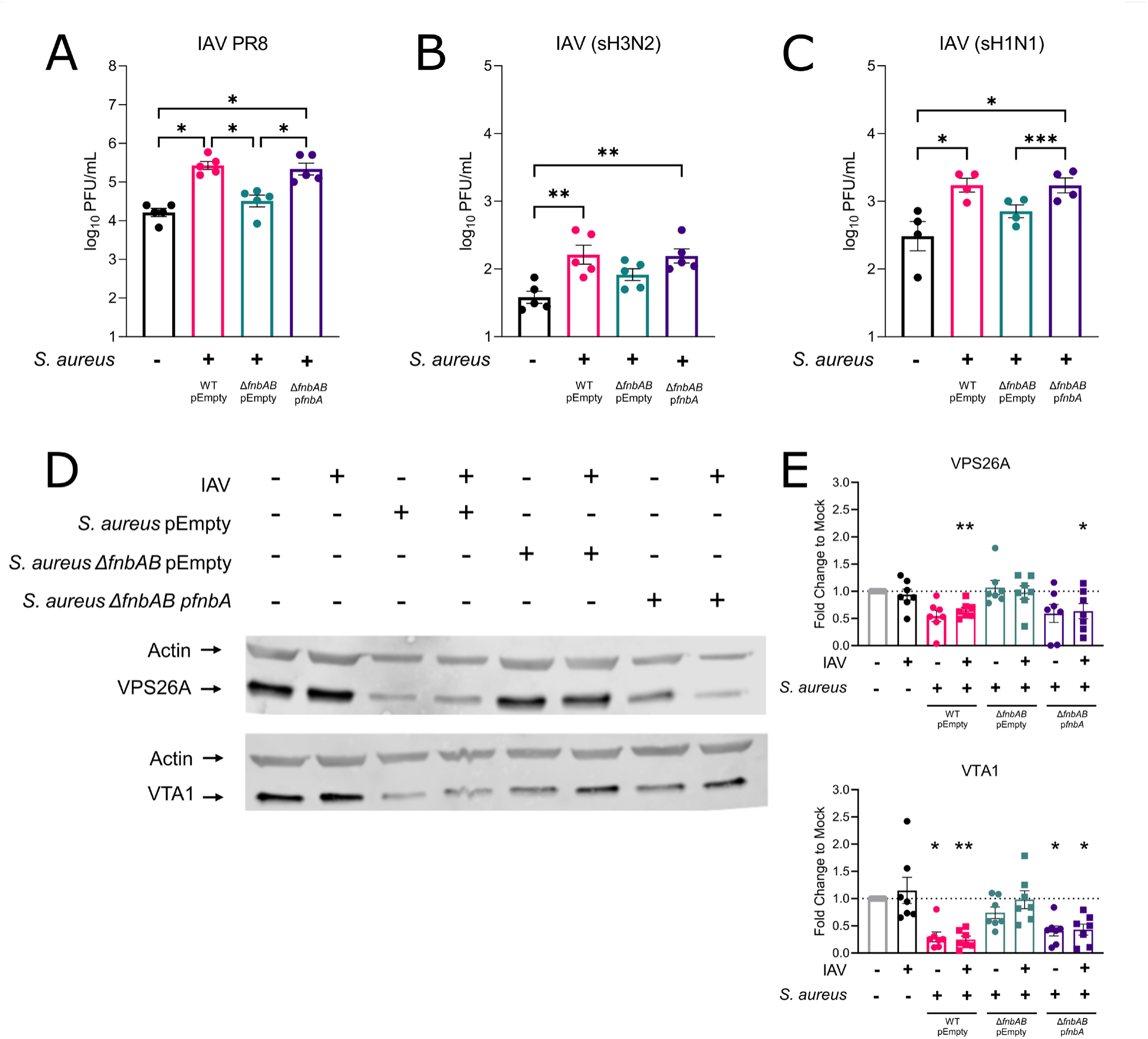
***S. aureus* enhances IAV replication through FnbA mediated decrease in VPS26A and VTA1.** Cells were infected as in Figure 1A, with **A)** A/Puerto Rico/8/34 (PR8), **B)** A//Baltimore/JH-335/2022 (sH3N2), or **C)** A/Baltimore/JH-22377/2022 (sH1N1), and WT or mutant *S. aureus* were added. At 12hpi the culture supernatant was centrifuged at 17,000 x *g* for 3 minutes and the resulting supernatant passed through a 13 mm, 0.22 µm filter. A standard plaque assay was then performed on MDCK cells. Data shown are mean ± SEM of 4-5 experiments. * is p value < 0.05, ** is p value < 0.01, *** is p value < 0.001 based on a repeated measures one-way ANOVA with Turkey’s multiple comparison correction. At 12hpi cells were harvested, lysed and Western blot performed for **D)** VPS26A and **F)** VTA1. Cellular actin was measured concurrently in each sample. **E)** and **G)** Analysis of western blot images was performed. For each image, values were normalized to actin and are shown as fold change of mock. For statistical analysis, actin normalized values were converted to log2, and a one-way ANOVA with repeated _measures_ was performed, with Turkey’s multiple comparison correction. ± SEM of 4-5 experiments. * is p value ≤ 0.05, ** is p value ≤ 0.01.

**Figure 5.**
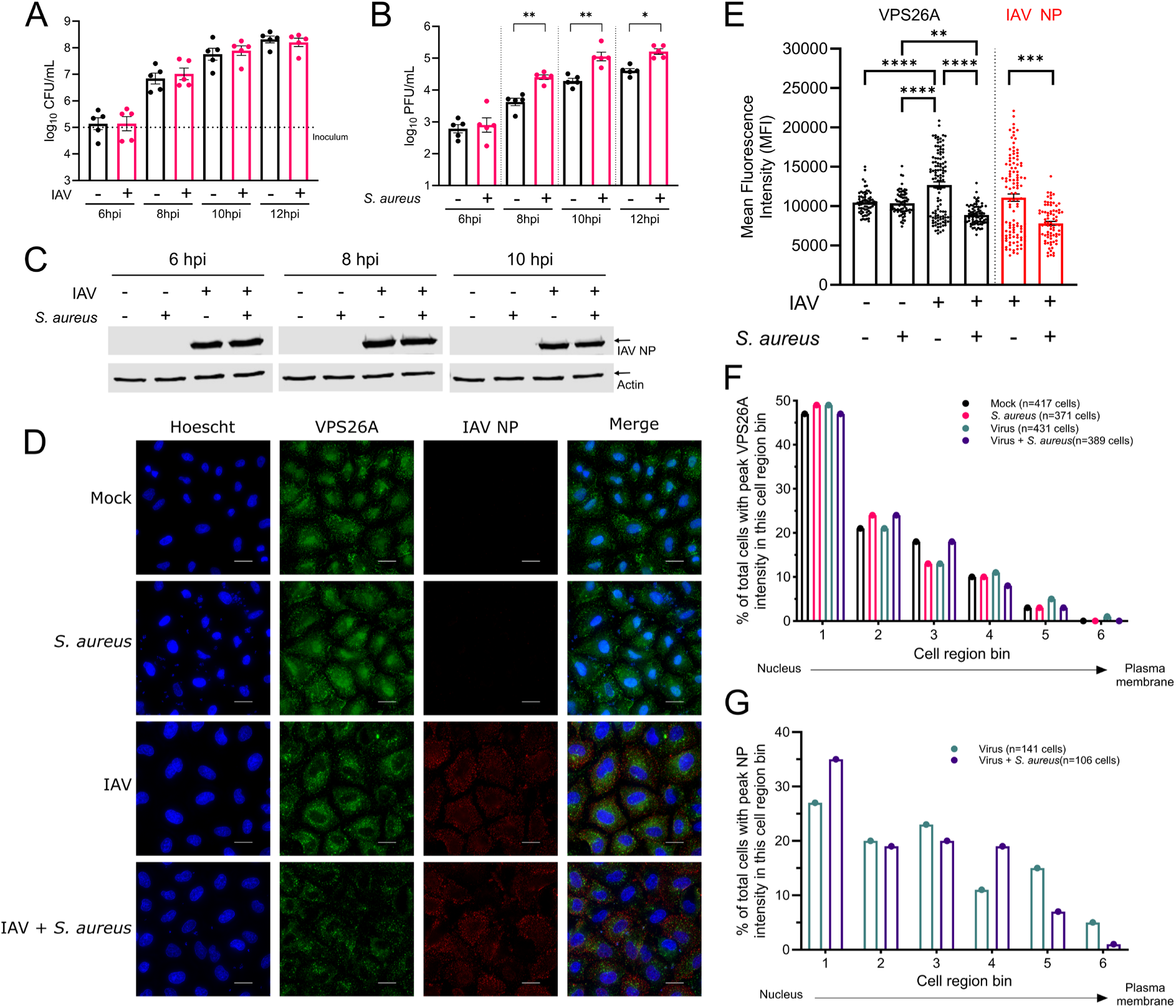
***S. aureus* co-infection enhances IAV within a single replication cycle.** Cells were infected as in Figure 1A. **A)** At the indicated times post infection (hpi), bacterial replication was determined by serial dilution in PBS and plating on TSA to calculate CFU/mL. Data shown are mean ± SEM of 5 experiments. **B)** Samples from **A)** were taken, centrifuged at 21,000 x *g* for 3 minutes and the resulting supernatant passed through a 13 mm, 0.22 µm filter. A standard plaque assay was then performed on MDCK cells. Infectious virus in the culture supernatant was quantified by plaque assay. * is p value ≤ 0.05, ** is p value ≤ 0.01, based on a two-tailed, paired student’s t test. **C)** Cells from A were lysed and used for a Western blot against IAV NP and β-actin. Representative images shown. **D)** Viral entry was synchronized by cold adsorption before addition of *S. aureus*. Cells were fixed at 8 hpi and stained for VPS26A and IAV NP. Representative immunofluorescence images. Scale bar is 10 µm. **E)** Mean fluorescence intensity of samples from **D)** Mean fluorescence intensity of VPS26A and NP was quantified per cell. Was quantified for VPS26A and NP. ** is p value < 0.01, *** is p< 0.001, **** is p< 0.0001, based on a one-way ANOVA with Turkey’s multiple comparison correction. Intracellular distribution of VPS26A **(F)** and NP (**G)** at 8hpi. Cells were divided into six equal regions relative to the nucleus, and each line profile was scored according to the region containing the highest fluorescence signal.

To determine whether the pro-viral effect was conserved across different IAV strains, we extended our analysis to seasonal isolates of H1N1 and H3N2 viruses. WT *S. aureus* significantly increased viral replication for both strains (Fig. 4B, 4C). Similar to our previous findings, the Δ*fnbAB* mutant lacked pro-viral activity against both seasonal isolates, while complementation with *fnbA* restored the phenotype (Fig. 4B, 4C).

Given that the changes in VPS26A and VTA1 were induced by *S. aureus*, we hypothesized that they were linked to the FnbA-dependent pro-viral phenotype. To test this, cells were infected as in Fig. 1A with WT, Δ*fnbAB*, or complemented *S. aureus*. At 12 hpi, VPS26A and VTA1 levels were assessed by Western blot. VPS26A levels were decreased in cells infected with WT *S. aureus*, but not the Δ*fnbAB* mutant, and this effect was restored by complementation with *fnbA* (Fig. 4 D, E). The same pattern was observed for VTA1 (Fig. 4 F, G). Together, these data demonstrate that FnbA mediates *S. aureus*-induced enhancement of IAV replication and concurrent changes in the abundance of host vesicular trafficking proteins.

### *S. aureus* enhancement of IAV occurs within a single replication cycle, without increasing viral protein synthesis

Having identified host and bacterial contributors to the pro-viral phenotype, we next sought to determine how these changes affect IAV replication. We first performed a time-course analysis to determine when the increase in infectious virus emerged. Bacterial replication increased from 6 hpi and plateaued between 10 and 12 hpi (Fig. 5A). The presence of *S. aureus* did not result in earlier production of infectious virus, but increased viral titres were detected by 8 hpi, coinciding with the first wave of progeny virus production (Fig. 5B). Peak viral titres occurred at 10 hpi both in the presence and absence of bacteria. Despite the increase in infectious virus, NP levels did not increase between IAV-infected and co-infected cells at any time point examined (Fig. 5C). Together, these data suggest that *S. aureus* enhances the production of infectious virus within a single IAV replication cycle without increasing viral protein abundance.

Given the role of VPS26A in host vesicular trafficking, we hypothesized that its reduction during co-infection could alter the intracellular trafficking of viral proteins to sites of assembly at the plasma membrane. To investigate this, we examined VPS26A together with the viral NP by immunofluorescence at 8 hpi. No clear co-localization between VPS26A and NP was observed (Fig. 5D). Consistent with our Western blot analysis, mean fluorescence intensity per cell confirmed reduced VPS26A abundance in the presence of *S. aureus*. In contrast, NP fluorescence intensity was not increased during co-infection and instead showed a modest decrease across the cell population (Fig. 5E).

We next examined whether co-infection altered the intracellular distribution of VPS26A or NP. Protein distribution relative to the nucleus was quantified by dividing individual cells into six equal regions (bins) and determining the region containing the highest fluorescence signal (Supplementary Fig 6). *S. aureus* induced no changes in the distribution of VPS26A (Fig 5F) and only a modest redistribution of NP was seen, with more cells displaying highest signal in intermediate regions of the cell (Fig 5G). Collectively, these data show *S. aureus* enhancement of IAV occurs withing a single replication cycle, with no increase in viral protein levels or changes in viral protein distribution.

**Figure 6.**
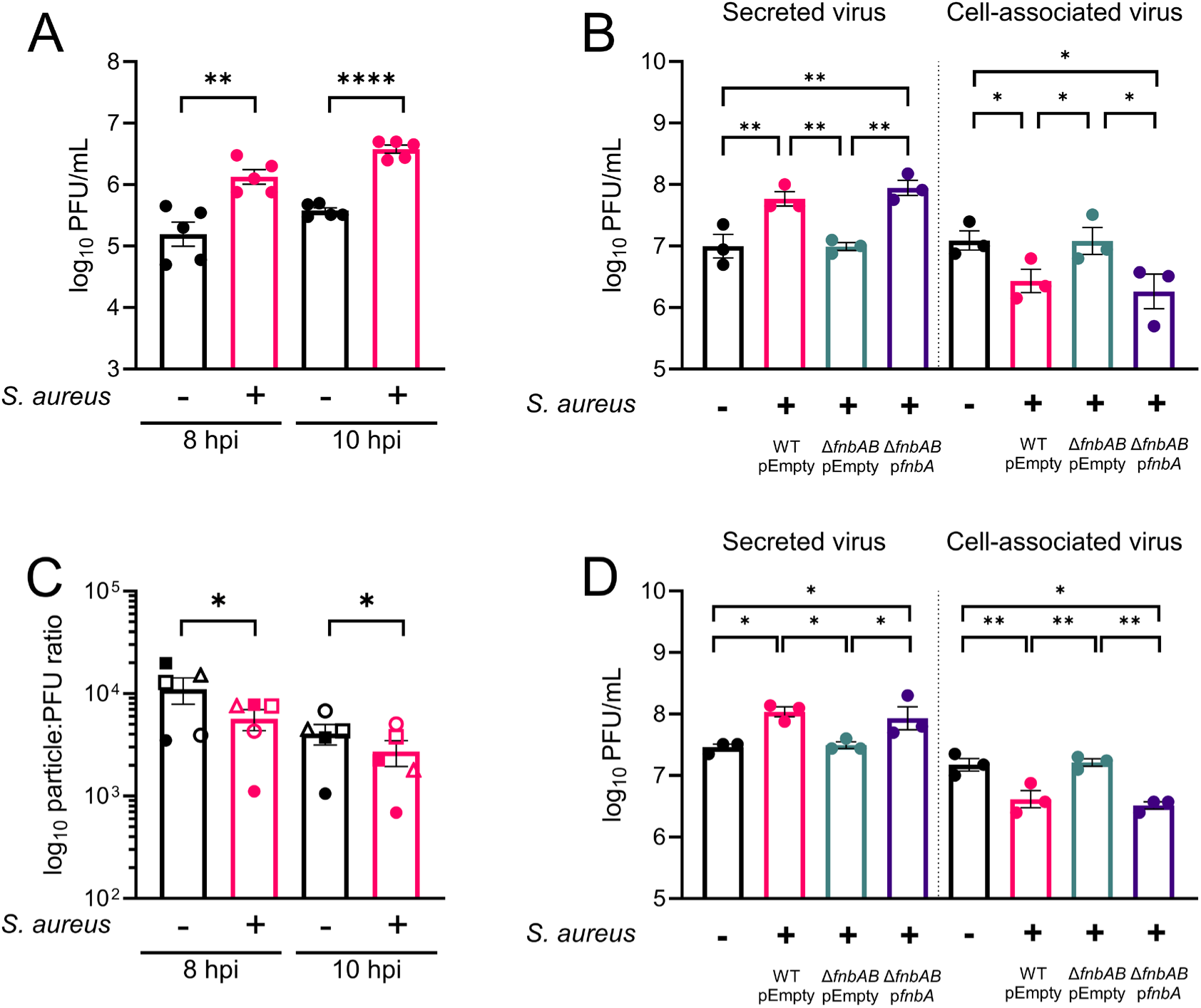
***S. aureus* enhances the formation and release of infectious virus particles during co-infection.** Cells were infected as in Fig 1A, except that viral entry was synchronized by cold adsorption. Samples were collected at 8 and 10 hpi. **A)** Infectious virus was quantified by plaque assay. Data shown are mean ± SEM of 5 experiments. ** is p value ≤ 0.01, **** is p value ≤ 0.0001 based on a two-tailed, paired student’s t test. **B)** Viral genome copies were quantified by RT-qPCR and particle-to-PFU ratios calculated relative to the corresponding infectious titre. Samples from the same experiment are denoted as matching datapoint symbols. * is p value ≤ 0.05, based on a two-tailed, lognormal paired student’s t test performed for each timepoint. Cells were infected as in Figure 1A with WT or mutant *S. aureus.* Samples were harvested at 8hpi (**C)** or 10 hpi **(D).** Infectious virus in the culture supernatant was quantified by plaque assay (left panels). Cells were collected by centrifugation and subjected to three freeze-thaw cycles before cell-associated infectious virus was quantified by plaque assay (right panels). Data shown are mean ± SEM of 3-5 experiments. * is p value ≤0.05, ** is p value ≤ 0.01 based on a repeated measures one-way ANOVA with Turkey’s multiple comparison correction.

### *S. aureus* co-infection enhances the production and release of infectious IAV particles

As increased infectious virus production occurred without corresponding changes in viral protein abundance or localization, we next examined the infectivity of the virus particles produced during co-infection. Infections were synchronized by cold adsorption, and particle-to-PFU ratios were determined at 8 and 10 hpi. Consistent with our previous findings, infectious viral titres were increased in the presence of *S. aureus* (Fig. 6A). Co-infection also significantly reduced the particle-to-PFU ratio (Fig. 6B), indicating that a greater proportion of the viral particles produced were infectious.

The magnitude of the reduction in particle-to-PFU ratio was smaller than the increase in infectious viral titre, suggesting that an additional mechanism may also contribute to the pro-viral phenotype. We therefore examined whether *S. aureus* also affected virion release. As observed in our previous experiments, co-infection increased viral titres in the culture supernatant (Fig. 6C, D). This was accompanied by a marked decrease in cell-associated virus (Fig. 6C, D), consistent with more efficient release of virions from infected cells. This effect was lost with the Δ*fnbAB* mutant and restored by complementation with *fnbA* (Fig. 6C, D). Together, these data demonstrate that *S. aureus* promotes the production and release of infectious IAV particles through an FnbA-dependent mechanism.

## Discussion

Bacterial co-infection is a major complication of influenza, but most studies have focused on how prior viral infection alters host immunity and increases susceptibility to subsequent bacterial disease. Much less is known about the reciprocal interaction and, in particular, whether specific bacterial factors directly alter IAV replication. We previously demonstrated that the secreted *S. aureus* Lipase 1 enhances IAV replication by modifying host lipids and promoting infectious virus production^20^. Here, we identify a second bacterial factor that directly promotes IAV replication and demonstrate that the *S. aureus* surface protein FnbA enhances infectious virus production across multiple IAV strains. This phenotype was associated with FnbA-dependent reductions in the host trafficking proteins VPS26A and VTA1. Although co-infection did not substantially alter viral protein abundance or intracellular distribution, it increased the proportion of infectious particles produced and promoted more efficient release of fully formed virions from infected cells. Together, these findings identify a previously unrecognized mechanism through which a bacterial surface protein can modify host cell biology to promote IAV replication.

Our identification of FnbA followed an unbiased screen of non-essential *S. aureus* genes and subsequent analysis of the SaeRS regulon. The involvement of SaeRS is consistent with previous work demonstrating that this regulatory system contributes to the severity of secondary *S. aureus* pneumonia following IAV infection^18^. Notably, the only strain in our initial comparison that lacked pro-viral activity was Newman, which carries a well-characterized mutation in *saeS* that results in altered SaeRS activity and regulation of its downstream targets^37,38^. Together, these findings support a role for SaeRS-regulated factors in enhancing IAV replication.

FnbA is a multifunctional cell wall-associated adhesin with established roles in *S. aureus* colonization and disease. Fibronectin-binding proteins promote adherence to human airway epithelial cells^41^ and contribute to invasive disease, including endocarditis and sepsis^40,42,43^. FnbA binds fibronectin, facilitating engagement of α5β1 integrin and bacterial internalization into non-phagocytic cells^38,44–46^. This interaction also activates host signalling pathways and induces cytoskeletal rearrangements. Our findings identify a new effect of FnbA on host cell biology, with FnbA mediating a decrease in VPS26A and VTA1. The mechanism connecting FnbA to these changes is not yet clear. Whether FnbA-mediated engagement of the fibronectin-integrin pathway contributes to the changes in VPS26A and VTA1 remains to be determined. Establishing this relationship will be important for understanding how FnbA alters host cell biology to promote IAV replication.

VPS26A is a component of the VPS26A-VPS29-VPS35 retromer complex, which controls sorting and recycling of cargo from endosomes^47,48^. Previous work has implicated this complex in IAV infection, specifically during viral entry. Poston *et al.* showed that loss of VPS29 enhanced IAV entry, with disruption of other retromer components producing a similar phenotype, supporting a role for the retromer complex in restricting an early stage of infection^49^. Here, we found that reduced VPS26A abundance was associated with increased infectious virus production at later stages of the replication cycle, extending the involvement of VPS26A beyond its previously described role during viral entry. This later effect may be particularly relevant given that retromer also contributes to intracellular trafficking of IAV M2, which undergoes retrograde transport through endosomes and the Golgi^50^. Determining how reduced VPS26A abundance reshapes retromer cargo trafficking may identify the cellular processes responsible for enhanced infectious virion production and release.

VTA1 provides a distinct potential link between altered host trafficking and IAV replication. VTA1 functions as a cofactor for the AAA ATPase VPS4, promoting VPS4 activity and ESCRT-III disassembly^23,51^. This relationship is particularly interesting in the context of IAV because, unlike many enveloped viruses, IAV does not require VPS4 or the canonical ESCRT machinery for efficient budding and release^25,52^. Instead, membrane scission during IAV budding can be mediated by the viral M2 protein^53^. Thus, the reduction in VTA1 observed during co-infection is unlikely to simply reflect disruption of ESCRT-dependent budding. Defining whether the effect of reduced VTA1 abundance on infectious virion production is dependent on VPS4 will distinguish between a canonical VTA1 function and an alternative, ESCRT-independent mechanism.

The increased production of infectious virus during co-infection appears to result from two complementary effects on progeny virions. Co-infection reduced the particle-to-PFU ratio, indicating that a greater proportion of the total particles produced were infectious. Concurrently, the extracellular infectious virus increased as the cell-associated fraction decreased. The latter phenotype was dependent on FnbA. These findings raise the possibility that the established interaction between FnbA and host fibronectin contributes to the increased release of infectious virus. Fibronectin has previously been implicated in IAV infection, with depletion or antibody-mediated inhibition of fibronectin reducing replication of human H1N1 viruses by interfering with post-attachment viral entry^54^. Fibronectin is also central to FnbA-mediated link between *S. aureus* and host cells, and both its abundance and organization can markedly influence these interactions^55^. Although these studies concern viral entry and bacterial internalization, respectively, they establish fibronectin as a shared host factor at the cell surface. It will therefore be interesting to determine whether FnbA alters the abundance or organization of fibronectin during co-infection and whether these changes reduce retention of newly formed IAV particles at the cell surface, thereby promoting their release.

Together, our findings demonstrate that *S. aureus* can enhance IAV replication through manipulation of host cell biology. We identify FnbA as a bacterial determinant of this interaction and link its pro-viral activity to reductions in VPS26A and VTA1 and increased production and release of infectious virions. FnbA is a highly multifunctional surface protein that binds several host extracellular matrix proteins, including fibronectin, fibrinogen and elastin^56,57^, facilitating bacterial adhesion and invasion and contributing to *S. aureus* pathogenesis. Despite decades of study, our findings identify an additional interaction between FnbA and the host, linking this protein to changes in cellular trafficking proteins and revealing consequences for the replication of a co-infecting virus. More broadly, this work demonstrates that bacteria can actively modify host processes to promote viral replication, highlighting the reciprocal nature of viral–bacterial interactions.

## Acknowledgements

This work was supported by Canadian Institutes for Health Research Project Grant 198010, a Natural Science and Engineering Research Council Discovery Grant RGPIN-2024-04587 and a BC Lung Foundation Research grant to MIG. We would like to thank Dr Stephen Tuffs for critical review of this manuscript.

