## Supplemental Information for "*Staphylococcus aureus* enhances influenza A virus replication through modulation of host cell vesicular transport"

Supplementary Table 1 – A list of bacterial strains used.

| **Species** | **Strain** | **Mutation** | **Plasmid** | **Source** |
|---|---|---|---|---|
| *Staphylococcus aureus* | USA300 | | | [17] |
| *Staphylococcus aureus* | USA300 | *ΔfnbAB* | | [17] |
| *Staphylococcus aureus* | USA300 | | pALC2073 | [17] |
| *Staphylococcus aureus* | USA300 | *ΔfnbAB* | pALC2073 | [17] |
| *Staphylococcus aureus* | USA300 | *ΔfnbAB* | pALC2073:*fnbA* | [17] |
| *Staphylococcus aureus* | USA300 | *saeR:tn* | | [18] |
| *Staphylococcus aureus* | USA300 | *saeR:tn* | pALC2073:*saePRQS* | This study |
| *Staphylococcus aureus* | USA300 | *saeS:tn* | | [18] |
| *Staphylococcus aureus* | USA300 | *saeS:tn* | pALC2073:*SaePRQS* | This study |
| *Staphylococcus aureus* | RN4220 | | | [17] |
| *Escherichia coli* | DH5α | | | Promega |
| *Staphylococcus aureus* | USA300 | *NTML collection* | | [18] |

Supplementary Table 2 – A list of primers used.

| Name | Sequence |
|---|---|
| Sae pALC F | GGGGGTACCTGAAGAAATGAGGAGTTAGCATG |
| Sae pALC R | GGGGAATTCCTCATTAATGGGAGCTTCTTT |
| IAV M qPCR F | GACCRATCCTGTCACCTCTGAC |
| IAV M qPCR R | AGGGCATTYTGGACAAAKCGTCTA |
| IAV Uni12 for vRNA | AGCAAAAGCAGG |

Supplementary Table 3 – *S. aureus* genes identified from genome wide screens to have significant differences in pro-viral activity.

| Protein name | Gene name | Accession number | Analyzed further | Reason for exclusion |
|---|---|---|---|---|
| hypothetical protein | | SAUSA300_0052 | No | Hypothetical |
| sucrose-specific PTS tranporter protein | | SAUSA300_0194 | | Growth phenotype |
| putative membrane protein | | SAUSA300_0230 | No | Growth phenotype, Hypothetical |
| putative membrane protein | | SAUSA300_0341 | No | Growth phenotype, Hypothetical |
| acetyltransferase | | SAUSA300_0451 | No | Growth phenotype |
| cobalamin synthesis protein/P47K family protein | | SAUSA300_0673 | | |
| anion transporter family protein | | SAUSA300_0676 | | |
| DNA-binding response regulator SaeR | *saeR* | SAUSA300_0691 | | |
| ABC transporter ATP-binding protein | | SAUSA300_0704 | | |
| hypothetical protein | | SAUSA300_0856 | No | Hypothetical |
| isopropylmalate synthase-related protein | | SAUSA300_0879 | No | Hypothetical |
| hypothetical protein | | SAUSA300_0903 | No | Hypothetical |
| methionine sulfoxide reductase A | *msrA* | SAUSA300_1256 | | |
| e ABC transporter, permease protein / oligopeptide permease, channel-forming | | SAUSA300_1276 | | |
| sensor histidine kinase protein | *arlS* | SAUSA300_1307 | | |
| putative 5'-3' exonuclease | | SAUSA300_1332 | No | Hypothetical |
| hypothetical protein | | SAUSA300_1336 | No | Growth phenotype, Hypothetical |
| phiSLT ORF 77-like protein | | SAUSA300_1415 | | |
| arginine repressor | *argR* | SAUSA300_1469 | | |
| phosphate starvation-induced protein, PhoH family | | SAUSA300_1531 | No | Growth phenotype |
| conserved hypothetical protein | | SAUSA300_1560 | No | Hypothetical |
| glutamyl-aminopeptidase | | SAUSA300_1691 | | |
| putative transposase | | SAUSA300_1732 | No | Hypothetical |
| conserved hypothetical protein | | SAUSA300_1853 | No | Hypothetical |
| two-component sensor histidine kinase | *VraS* | SAUSA300_1866 | | |
| hypothetical protein | | SAUSA300_1868 | No | Growth phenotype, Hypothetical |
| exonuclease | | SAUSA300_1875 | | |
| high affinity proline permease | *putP* | SAUSA300_1883 | | |
| isopropylmalate isomerase small subunit | *leuD* | SAUSA300_2013 | | |
| MarR family transcriptional regulator | | SAUSA300_2216 | Yes | Growth phenotype |
| urease subunit beta | *ureB* | SAUSA300_2239 | | |
| conserved hypothetical protein | | SAUSA300_2528 | Yes | Hypothetical |
| putative glycosyl transferase | | SAUSA300_2583 | Yes | Growth phenotype, Hypothetical |

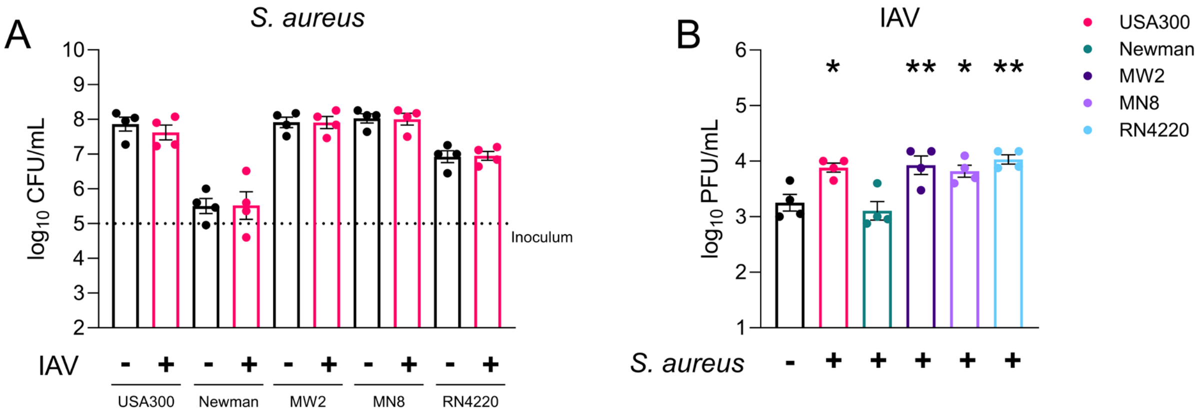


**Supplementary Figure 1 – *S. aureus* pro-viral activity is conserved across most laboratory strains**. A549 cells in a 12 well plate were infected with A/Puerto Rico/8/34 at MOI 3 for 1h. Inoculum was removed, extracellular virus lysed with an acid wash for 90 sec, and fresh media added. $1x10^5$ CFU/mL of *S. aureus* were added to the extracellular environment. **A)** At 12h post infection (hpi) a sample from the supernatant was taken, serially diluted in PBS, plated on TSA plates and incubated at 37°C for 18h. The dotted inoculum line represents the number of bacteria added at 0h. **B)** Samples from **A)** were collected, centrifuged at 17,000 x *g* for 3 minutes and the resulting supernatant passed through a 13 mm, 0.22 µm filter. A standard plaque assay was then performed on MDCK cells to determine infectious virus titre. Data shown are mean ± SEM of 4 experiments. * is p value ≤ 0.05, based on a one-way ANOVA with Dunnett's multiple comparison correction.

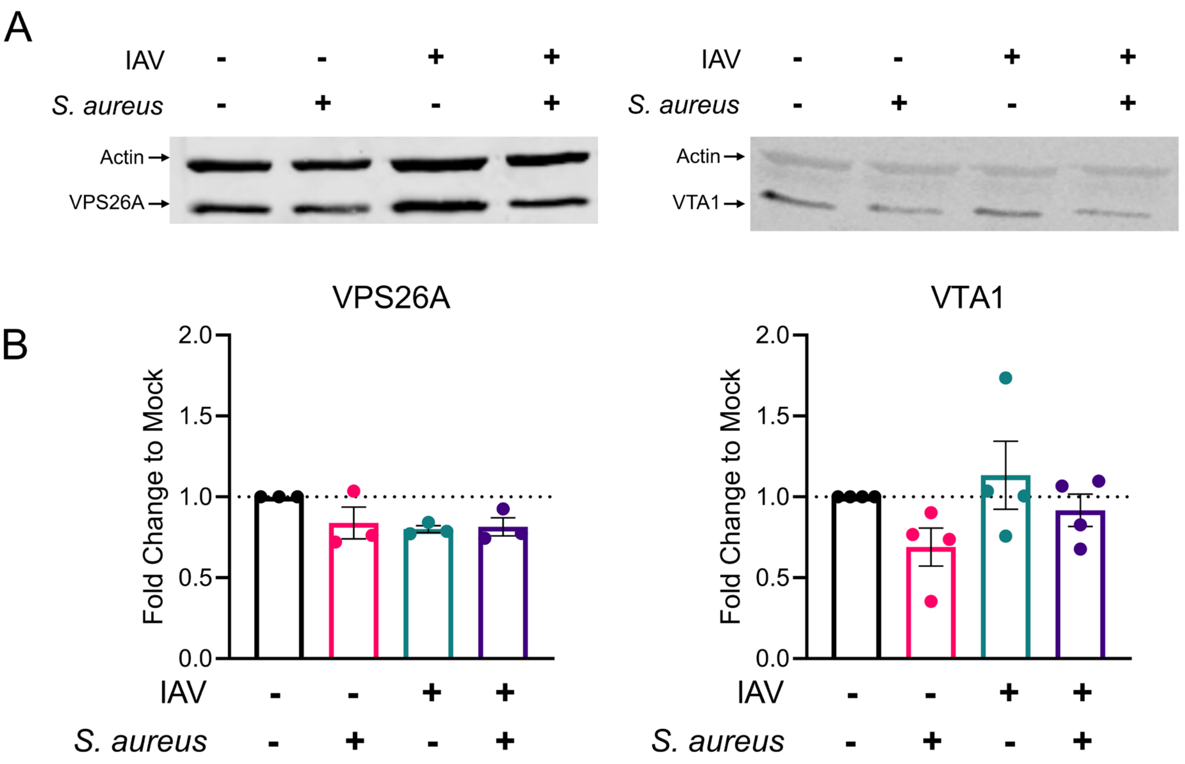


**Figure 2 – *S. aureus* co-infection triggers changes in host cell trafficking proteins.** Cells were infected as in Figure 1A. At 12hpi cells were harvested, lysed and Western blot performed for **A)** VPS26A and **B)** VTA1. Cellular actin was measured concurrently in each sample. **C)** and **D)** Analysis of western blot images was performed. For each image, values were normalized to actin and are shown as fold change of mock.

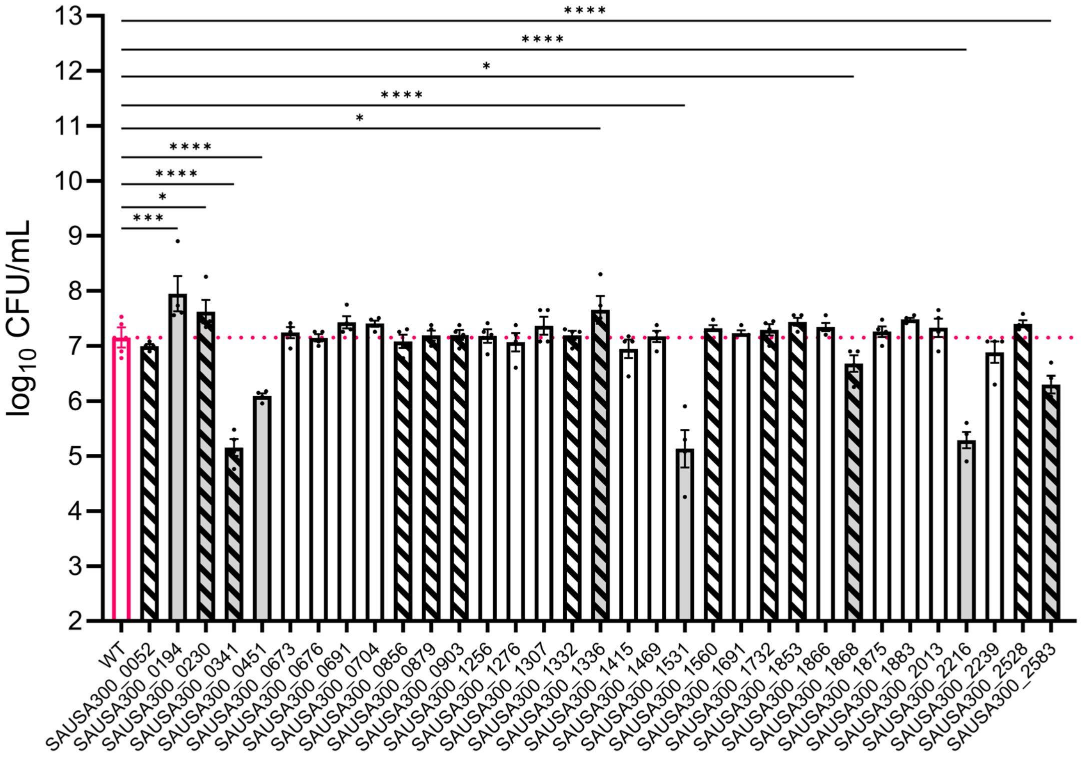


**Figure 3. *S. aureus* growth on A549 cells is affected by some gene mutations.** A549 cells in a 96 well plate were washed 3 times, and serum-free media added. ~$5 \times 10^4$ CFU of WT or mutant *S. aureus* were added to the media. At 12hpi a sample from the supernatant was taken, serially diluted in PBS, plated on TSA plates and incubated at 37°C for 18h. The pink dotted line represents the average value of the WT. Bars with grey shading indicate mutants that show statistically significant difference in growth compared to the WT. Bars with black pattern fills indicate genes that are not known or characterized. Data shown are mean ± SEM of 4 experiments. * is p value ≤ 0.05, *** is p ≤ 0.001, **** is ≤ 0.0001, based on a one-way ANOVA with Fisher's exact LSD test.

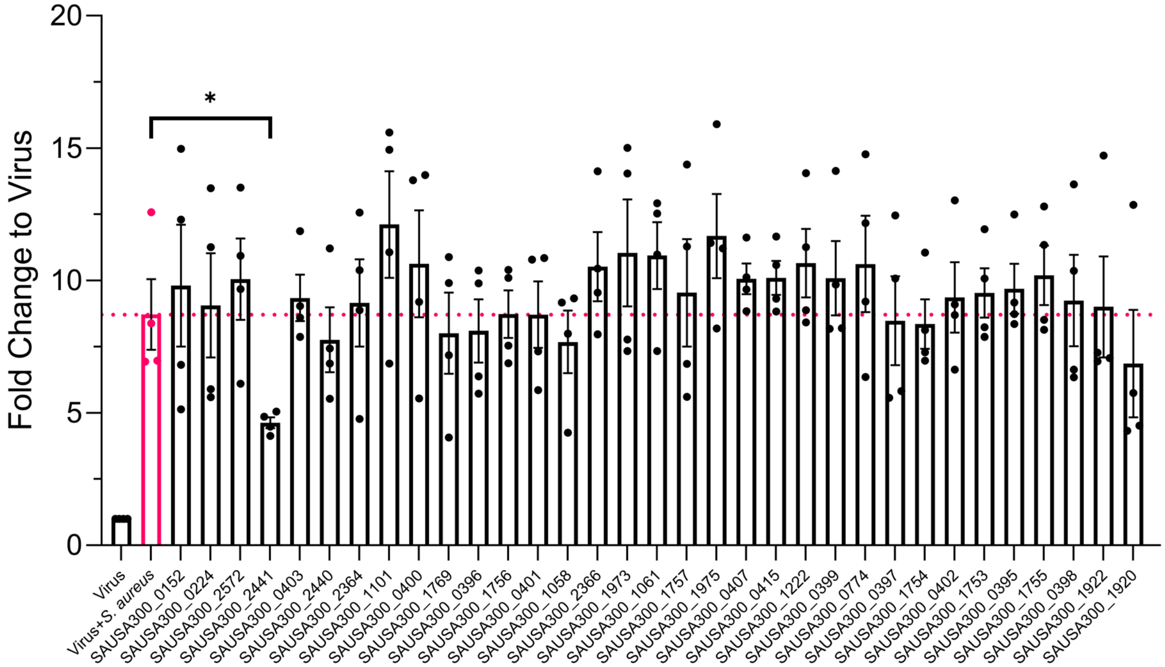


**Supplementary Figure 4 – *S. aureus* pro-viral activity is eliminated when fnbA is mutated.** A549 cells in a 96 well plate were infected with IAV WSN-Luc at a MOI of 2 for 1h. Inoculum was removed and serum-free media added. ~5x$10^4$ CFU of *S. aureus* WT or mutant bacteria were added to the media. At 12hpi a sample from the supernatant was taken, mixed with equal volume of Nano-Glo reagent and luciferase measured on a plate reader. All values were normalized to virus only samples. The pink dotted line represents the average value of the WT. Data shown are mean ± SEM of 4 experiments. * is p value ≤ 0.05, based on a one-way ANOVA with Fisher's exact LSD test.

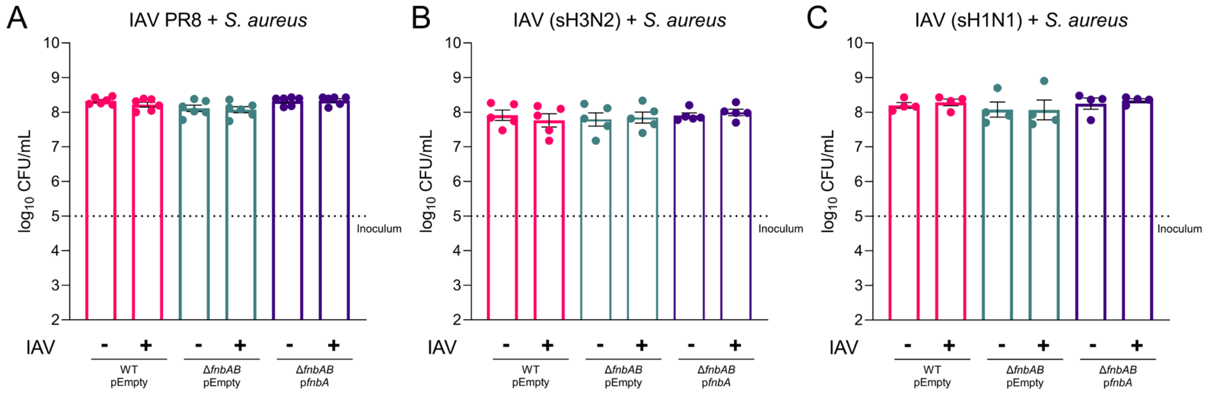


**Supplementary Figure 5 – *S. aureus* growth is not impacted by prior infection with IAV.** A549 cells were infected with IAV A/Puerto Rico/8/34 (PR8) **(A)**, seasonal IAV H1N1 **(B)** or seasonal IAV H3N2 **(C)** at a MOI of 3 for 1h. Inoculum was removed and serum-free media added. 1x10$^{5}$ CFU of *S. aureus* strains indicated were added to the media. At 12hpi a sample from the supernatant was taken, serially diluted in PBS, plated on TSA plates and incubated at 37°C for 18h. The dotted inoculum line represents the number of bacteria added at 0h. Data shown are mean ± SEM of 4-6 experiments.

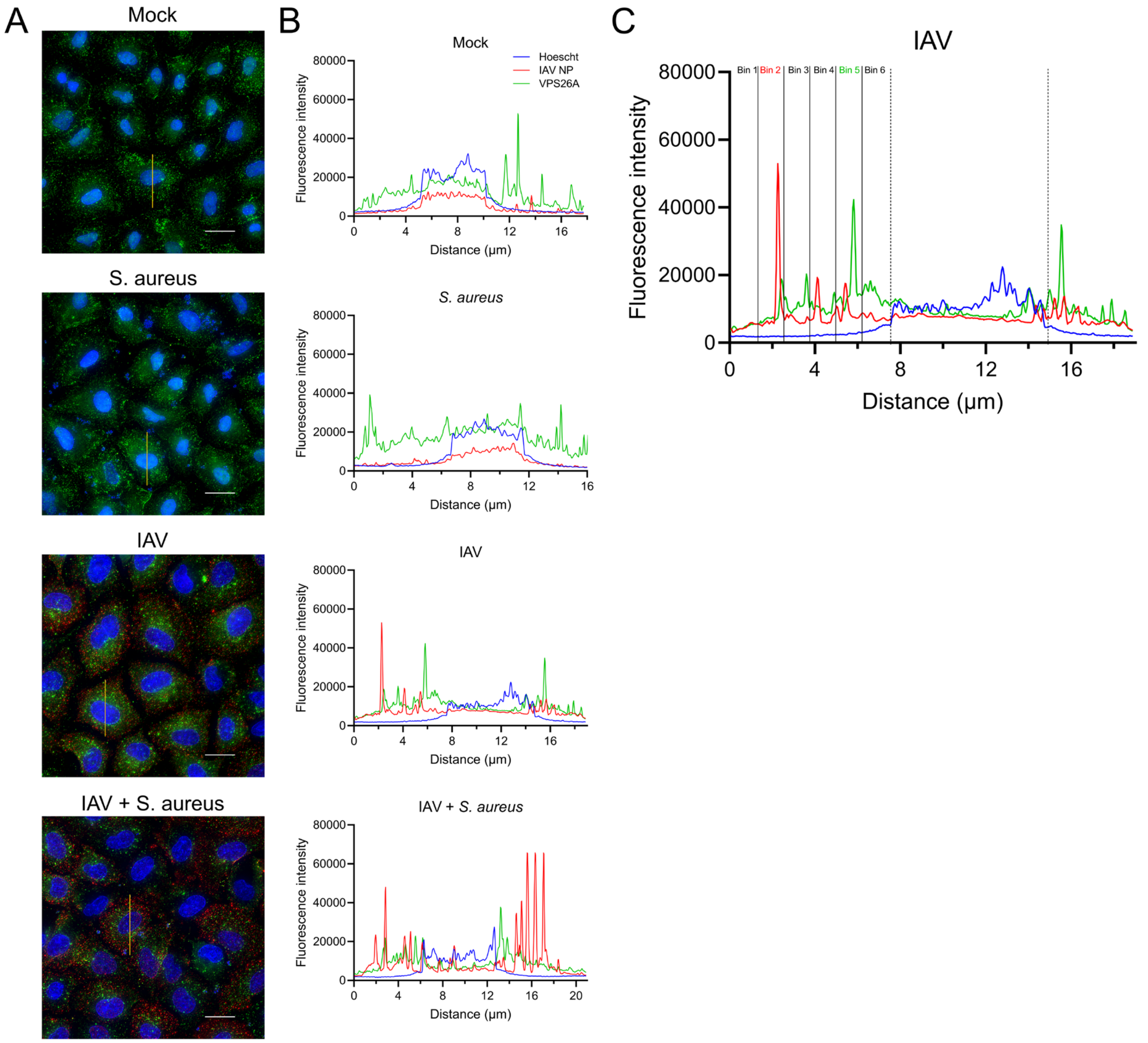


**Figure 6 – Method of quantification of fluorescence intensity for the separation of cells into distinct regions (bins).** Cells were infected as in Figure 1A, except viral entry was synchronized by cold adsorption before addition of *S. aureus*. Cells were fixed at 8 hpi and stained for VPS26A and IAV NP. **A)** Representative immunofluorescence images of merged channels. The yellow line shows the line graph taken for this specific cell. **B)** Fluorescence intensity over the line graph from **A)**. **C)** Schematic representation of the separation of cell regions into bins. To account for differences in cell size, each line profile was divided into six equal sections spanning the cell region outside of the nucleus. Fluorescence within each section was quantified by calculating the area under the curve of the intensity profile. Each line profile was then assigned a score from 1 to 6 corresponding to the section with the highest signal from area under the curve. In this image highest signal for NP (in red) is in Bin 2, which is shown in red text; highest signal for VPS26A (in green) is in Bin 5, which is shown in green text.
